# Aquaglyceroporin AQP7’s affinity for its substrate glycerol

**DOI:** 10.1101/2021.11.23.469753

**Authors:** Michael Falato, Ruth Chan, Liao Y. Chen

## Abstract

AQP7 is one of the four human aquaglyceroporins that facilitate glycerol transport across the cell membrane, a biophysical process that is essential in human physiology. Therefore, it is interesting to compute AQP7’s affinity for its substrate (glycerol) with reasonable certainty to compare with the experimental data suggesting high affinity in contrast with most computational studies predicting low affinity. In this study aimed at computing the AQP7-glycerol affinity with high confidence, we implemented a direct computation of the affinity from unbiased equilibrium molecular dynamics (MD) simulations of three all-atom systems constituted with 0.16M, 4.32M, and 10.23M atoms, respectively. These three sets of simulations manifested a fundamental physics law that the intrinsic fluctuations of pressure in a system are inversely proportional to the system size (the number of atoms in it). These simulations showed that the computed values of glycerol-AQP7 affinity are dependent upon the system size (the inverse affinity estimations were, respectively, 47.3 mM, 1.6 mM, and 0.92 mM for the three model systems). In this, we obtained a lower bound for the AQP7-glycerol affinity (an upper bound for the dissociation constant). Namely, the AQP7-glycerol affinity is stronger than 1087/M (the dissociation constant is less than 0.92 mM). Additionally, we conducted hyper steered MD (hSMD) simulations to map out the Gibbs free-energy profile. From the free-energy profile, we produced an independent computation of the AQP7-glycerol dissociation constant being approximately 0.18 mM.

**Table of contents entry:** 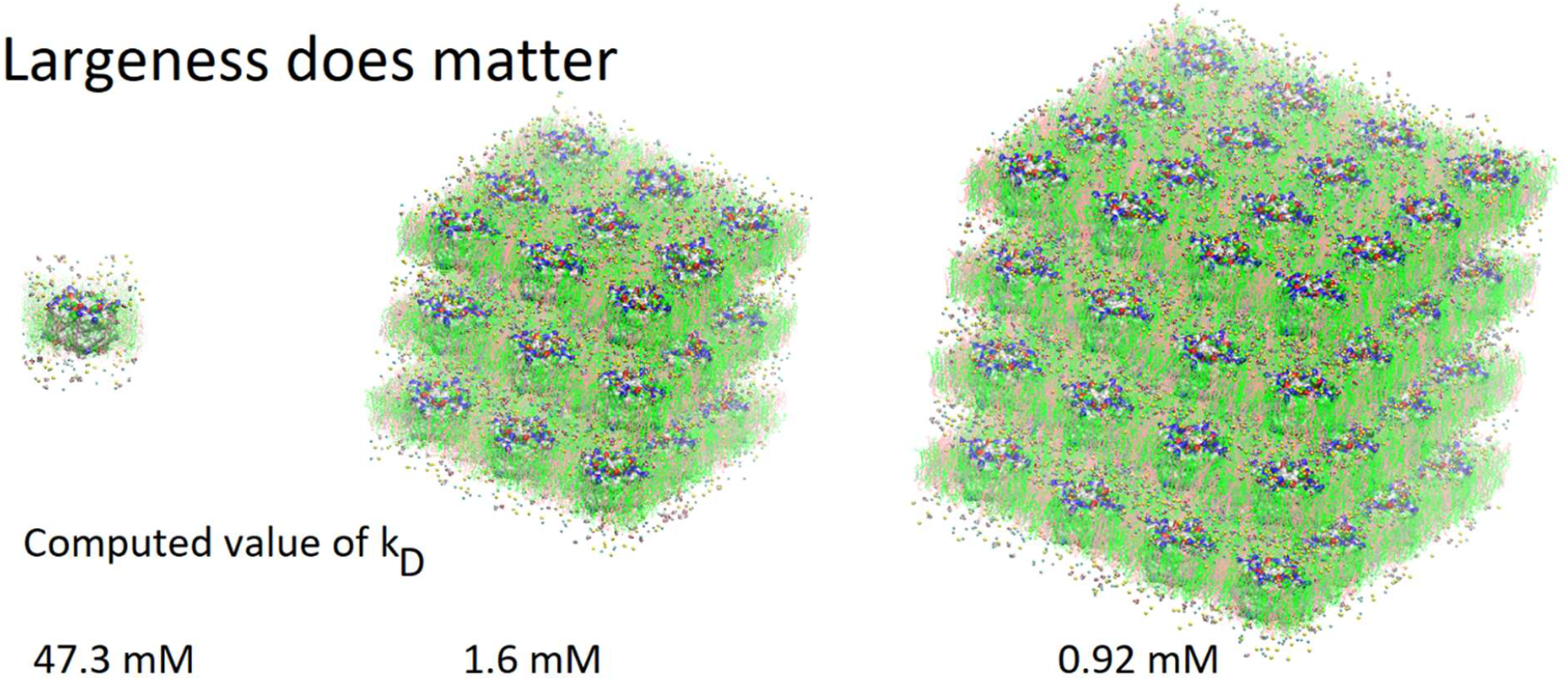

## INTRODUCTION

Aquaglyceroporins (AQGPs) are a subfamily of aquaporin (AQP) proteins[1-3] responsible for facilitated diffusion of glycerol and some other small neutral solutes across the cell membrane along the solute concentration gradient[4]. They also conduct water transport down the osmotic gradient. Among the 13 human AQPs, the AQGP subfamily consists of AQPs 3, 7, 9, and 10. AQGPs are fundamental to many physiological processes. For example, pancreatic AQP7 is involved in insulin secretion; all AQGPs participate in fat metabolism. Therefore, AQGPs are investigated as drug targets for metabolic diseases[5].

Among the many experimental and theoretical-computational investigations of aquaglyceroporins, one fundamental question remains: Does an AQGP have affinity for its substrate glycerol? In functional characterization experiments in 1994[6], *Escherichia coli* aquaglyceroporin GlpF was shown to facilitate unsaturable uptake of glycerol up to 200 mM into Xenopus oocytes, suggesting that GlpF has very low affinity for its substrate glycerol. In a series of functional experiments from 2008 to 2014[7-9], human aquaglyceroporins AQP7, AQP9, and AQP10 were shown to conduct saturated transport of glycerol with Michaelis constants around 10 µM, indicating that human AQGPs have high affinities for glycerol. In the crystal structures available to date (GlpF in 2000[10], *Plasmodium falciparum* PfAQP in 2008[11], AQP10 in 2018[12], and AQP7 in 2020[13-15]), glycerol molecules were found inside the AQGP channel and near the channel openings on both the intracellular (IC) and the extracellular (EC) sides, showing that all four AQGPs have affinities for glycerol. If we insisted that unsaturated transport precludes high affinity, these experimental data would suggest inconsistency. However, in an *in silico-in vitro* study[16] of glycerol uptake into human erythrocytes through AQP3[17], it was shown that an AQGP (having high affinity for its substrate glycerol) can conduct glycerol transport that is unsaturated up to 400 mM. The transport pathway for unsaturated transport through a high affinity facilitator protein was shown to involve two glycerol molecules next to each other both bound inside an AQP3 channel (one at the high affinity site and one at a low affinity site) for the transport of one glycerol molecule across the cell membrane[16]. It is the substrate-substrate interactions (mostly repulsion due to steric exclusion) inside a single-file channel that make it easy for two glycerol molecules cooperatively to move one substrate molecule across the AQGP channel *via* the high affinity site.

On the theoretical-computational side, the predicted affinities of AQGPs were derived from the computed free-energy profiles---the PMF curves (the potential of mean force as a function of an order parameter, namely, the Gibbs free energy of the system when the chosen degrees of freedom are set to a given set of values). The predictions are dependent upon the methods of computation used in a given study. For example, the estimated values of the glycerol-GlpF affinity range from < 1/*M* (from the PMF curve of Refs. [18, 19]) to > 10^3^/*M* (from the PMF curve of Ref. [20]). Currently, the estimations of the glycerol-AQP7 affinity stand at < 1/*M* (from the PMF curves of Refs. [13, 15]) in contrast with the experimental data of Ref. [7] showing high affinity ∼10^5^/*M*. All these point to the need of further theoretical-computational studies of AQGP-glycerol affinities.

In this research, we aim to reach computational convergence on glycerol-AQP7 affinity. We first carried out direct computations of the glycerol-AQP7 affinity by estimating the probability *p*_b_ of glycerol binding inside an AQP7 channel for a given glycerol concentration *c*_G_. The dissociation constant *k*_D_ = *c*_G_(1 − *p*_b_)/*p*_b_ is the inverse of the glycerol-AQP7 affinity. Running equilibrium molecular dynamics (MD) without any biases or constraints on three systems ranging from approximately 0.2 M atoms to 10 M atoms in sizes, we observed a convergence toward high AQP7-glycerol affinity. We also examined the intrinsic fluctuations of the model systems (Fig. 1). We found that the pressure fluctuations are inversely proportional to the system size as expected based on statistical thermodynamics[21]. In a system (consisting of 0.2 M atoms) typical in the current literature, the room mean squared pressure fluctuations >100 bar in the simulation of a system under a constant pressure of 1.0 bar. In another word, the model system is subject to constant agitations of an artificial sonicator in inverse proportion to the system size. These agitations are expected to loosen the binding between a protein and its substrate and thus to reduce the apparent affinity (*i*.*e*., the computed value of the glycerol-AQGP affinity). Our simulations of various system sizes showed that the computed values of glycerol-AQP7 affinity are strongly dependent upon the system size and that convergence of computational studies points to strong affinity between an AQGP and its substrate instead of weak affinity observed in small simulations. Seeking an independent confirmation of strong AQGP-glycerol affinity, we also determined the glycerol-AQP7 affinity from the PMF curve that was computed from a large set of hyper steered MD (hSMD) simulations.

**Fig 1.**
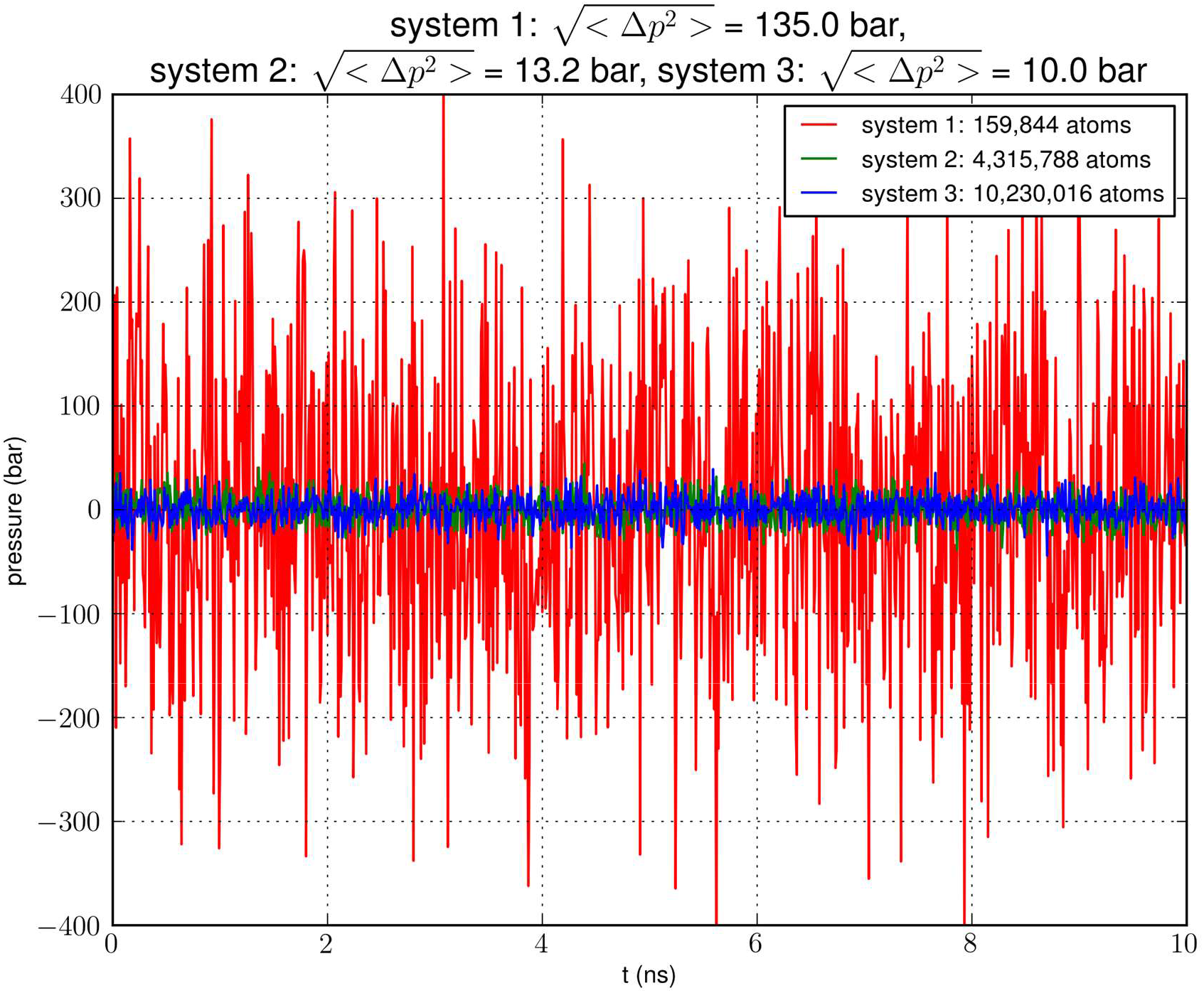
Pressure fluctuation in an NPT simulation of a small system (159,844 atoms) containing a single AQP7 tetramer, a large system (4,315,788 atoms) with 27 AQP7 tetramers, or a huge system (10,230,016 atoms) with 64 AQP7 tetramers. The pressure fluctuation is approximately proportional to the inverse of the system size in terms of atom numbers.

## METHODS

The parameters, the coordinates, and the scripts for setting up the model systems, running the simulations, and analyzing the data are available at Harvard Dataverse[22].

### Model system setup and simulation parameters

Following the well-tested steps in the literature, we employed CHARMM-GUI[23-25] to build an all-atom model of an AQP7 tetramer embedded in a 117Å×117Å patch of membrane (lipid bilayer). The AQP7 coordinates were taken from Ref. [13] (PDB: 6QZI). The positioning of the AQP7 tetramer was determined by matching the hydrophobic side surface with the lipid tails and aligning the channel axes perpendicular to the membrane. The AQP7-membrane complex was sandwiched between two layers of TIP3P waters, each of which was approximately 30Å thick. The system was then neutralized and salinated with Na^+^ and Cl^−^ ions to a salt concentration of 150 mM. Glycerol was added to the system to 50 mM in concentration. The system so constructed consists of a single AQP7 tetramer (four monomer channels) constituted with 159,844 atoms, which is referred to as SysI (shown in Supplemental Information, SI, Fig. S1). We employed NAMD 2.13 and 3.0 [26, 27] as the MD engines. We used CHARMM36 parameters[28-30] for inter- and intra-molecular interactions. We followed the literature’s standard steps to equilibrate the system[15, 31-33]. Then we ran unbiased MD for 2,000 ns (namely, 8,000 monomer·ns) with constant pressure at 1.0 bar (Nose-Hoover barostat) and constant temperature at 303.15 K (Langevin thermostat). The Langevin damping coefficient was chosen to be 1/ps. The periodic boundary conditions were applied to all three dimensions. The particle mesh Ewald (PME) was used for the long-range electrostatic interactions (grid level: 128×128×128). The time step was 2.0 fs. The cut-off for long-range interactions was set to 10 Å with a switching distance of 9 Å. The last 500 ns (2,000 monomer·ns) of the trajectory was used in the computation of the glycerol-AQP7 affinity.

We replicated SysI 26 times to obtain 27 copies of SysI. With appropriate translations of these copies, we formed SysII consisting of 27 AQP7 tetramers (illustrated in SI, Fig. S2). Unbiased MD was run for 15,000 monomer·ns for this large SysII with identical parameters used for SysI except that the PME was implemented on a grid of 384×384×384. The last 5,000 monomer·ns were used in the computation of the glycerol-AQP7 affinity. Likewise, we replicated SysI 63 times to form SysIII consisting of 64 AQP7 tetramers (illustrated in SI, Fig. S3). We ran unbiased MD on SysIII (with PME grid of 512×512×512) for 15,000 monomer·ns and used the last 5,000 monomer·ns in the computation of the glycerol-AQP7 affinity.

### Direct computation of AQP7-glycerol affinity

We used the part of an MD trajectory when the system is fully equilibrated to compute the probability *p*_b_ for an AQP7 channel being occupied with a glycerol molecule (being inside the single-file region of the channel, 7.1 Å to the IC/EC side from the NAA/NPS motifs illustrated in Fig. 2). Based on the equilibrium kinetics, *p*_b_ = *c*_G_/(*c*_G_ + *k*_D_) with *c*_G_ being the glycerol concentration, we computed the dissociation constant from the binding probability: *k*_D_ = *c*_G_(1 − *p*_b_)/*p*_b_.

**Fig 2.**
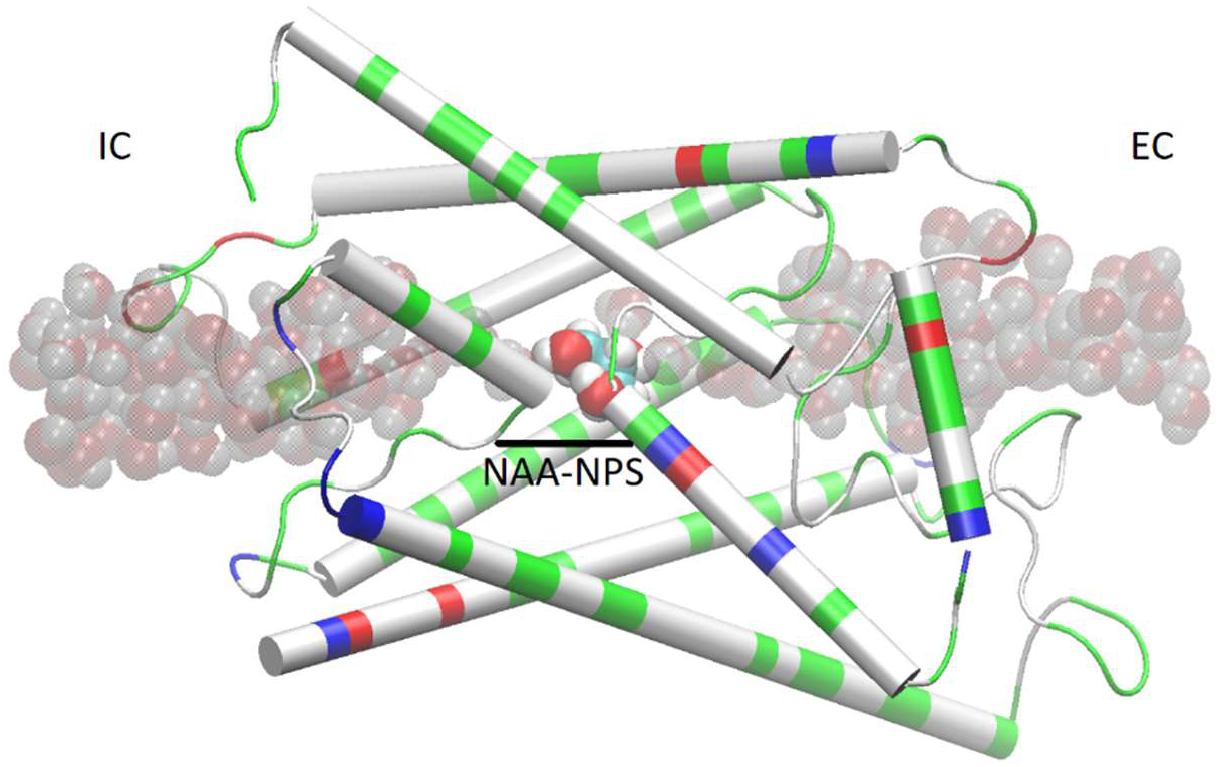
AQP7 monomer channel with a glycerol molecule (large spheres colored by atoms: C, cyan; O, red; H, white) at the central binding site near the NAA/NPS motifs. The whole monomer protein is shown as cartoons colored by residue types (positively charged, blue; negatively charged, red; hydrophilic, green; hydrophobic, white). The water molecules inside and near the channel are shown in shadowy spheres colored by atoms (O, red; H, white). All molecular graphics in this paper were rendered with VMD[35].

### Computing the Gibbs free-energy profile and the affinity

We conducted 2,100 ns hSMD of SysI (illustrated in Fig. 2) to compute the PMF along the glycerol transport path through an AQP7 channel across the membrane. We followed the multi-sectional protocol detailed in Ref.[34]. We defined the forward direction as along the z-axis pointing from the intracellular side to the extracellular side. We divided the entire glycerol transport path across the membrane from *z* = −28Å to *z* = 22Å into 50 evenly divided sections. From the central binding site (*z* = −1Å, shown in Fig. 2) to the EC side (*z* ≥ 22Å), the center-of-mass z-degree of freedom of glycerol was steered at a speed of 0.25 Å/ns for 4 ns over one section for a z-displacement of 1.0 Å to sample a forward path over that section. At the end of each section, the z-coordinate of the glycerol center-of-mass was fixed (or, technically, pulled at a speed of 0.0 Å/ns) while the system was equilibrated for 10 ns. From the end of the 10 ns equilibration, the z-coordinate of the glycerol center-of-mass was pulled for 4 ns for a z-displacement of −1.0 Å to sample a reverse path. From the binding site (*z* = −1Å) to the IC side (*z* ≤ −28Å), the center-of-mass z-degree of freedom of glycerol was steered for 4 ns for a z-displacement of −1.0 Å to sample a reverse path over one section. At the end of that section, the z-coordinate of the glycerol center-of-mass was fixed while the system was equilibrated for 10 ns. From the end of the 10 ns equilibration, the z-coordinate of the glycerol center-of-mass was pulled for 4 ns for a z-displacement of +1.0 Å to sample a forward path. In this way, section by section, we sampled a set of four forward paths and four reverse paths in each of the 50 sections (28 sections from the central binding site to the IC side and 22 sections from the central binding site to the EC side) along the entire transport path between the IC and the EC sides. The force acting on the glycerol center-of-mass was recorded along the forward and the reverse pulling paths for computing the PMF along the entire transport path from the IC side to the central binding site and then to the EC side. The PMF was computed from the work along the forward paths and the work along the reverse paths (SI, Fig. S6) *via* the Brownian-dynamics fluctuation-dissipation theorem[34].

Following the standard literature (*e*.*g*., [36]), one can relate the binding affinity (inverse of the dissociation constant k_Di_) at the *i*-th binding site to the PMF difference in 3 dimensions (3D) and the two partial partitions as follows:

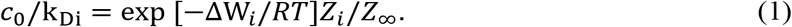

Here Δ*W*_i_ is the PMF at the *i*-th binding site minus the PMF in the dissociated state when glycerol is far away from the protein. *R* is the gas constant. *T* is the absolute temperature. *Z*_i_ is the partial partition of glycerol in the *i*-th bound state which can be computed by sampling the fluctuations in 3 degrees of freedom of the glycerol center of mass and invoking the Gaussian approximation for the fluctuations in the bound state[37]. *Z*_∞_ = 1/*c*_0_ is the corresponding partial partition in the dissociated state with *c*_0_ = 1*M* being the standard concentration.

## RESULTS AND DISCUSSION

### The long process of equilibration in a glycerol-AQP7 system

We conducted an MD run (under constant temperature and constant pressure, NPT) for 2,000 ns to fully equilibrate SysI consisting of one AQP7 tetramer (four AQP7 monomer channels) constituted with 159,844 atoms (SI, Fig. S1). We computed the root mean squared deviation (RMSD) from the crystal structure for each of the four protein monomers, which are shown in Fig. 3. We learnt that equilibrium was not reached until after 1,500 ns (*i*.*e*., 6,000 monomer•ns). Only during the last 500 ns, we observed multiple events of glycerol moving into and out of the AQP7 channel as illustrated in Movie 1. Therefore, only the last 500 ns (2,000 monomer•ns) of the MD trajectory should be used in the statistical analyses of the system. The earlier part of the trajectory represents a transient process toward equilibrium which is inevitably dependent upon the initial conditions of the model system. Multiple monomer·µs simulations are necessary for significant sampling of glycerol-AQP7 kinetics. This indicates that glycerol-AQP7 interactions are strong rather than weak.

**Fig 3.**
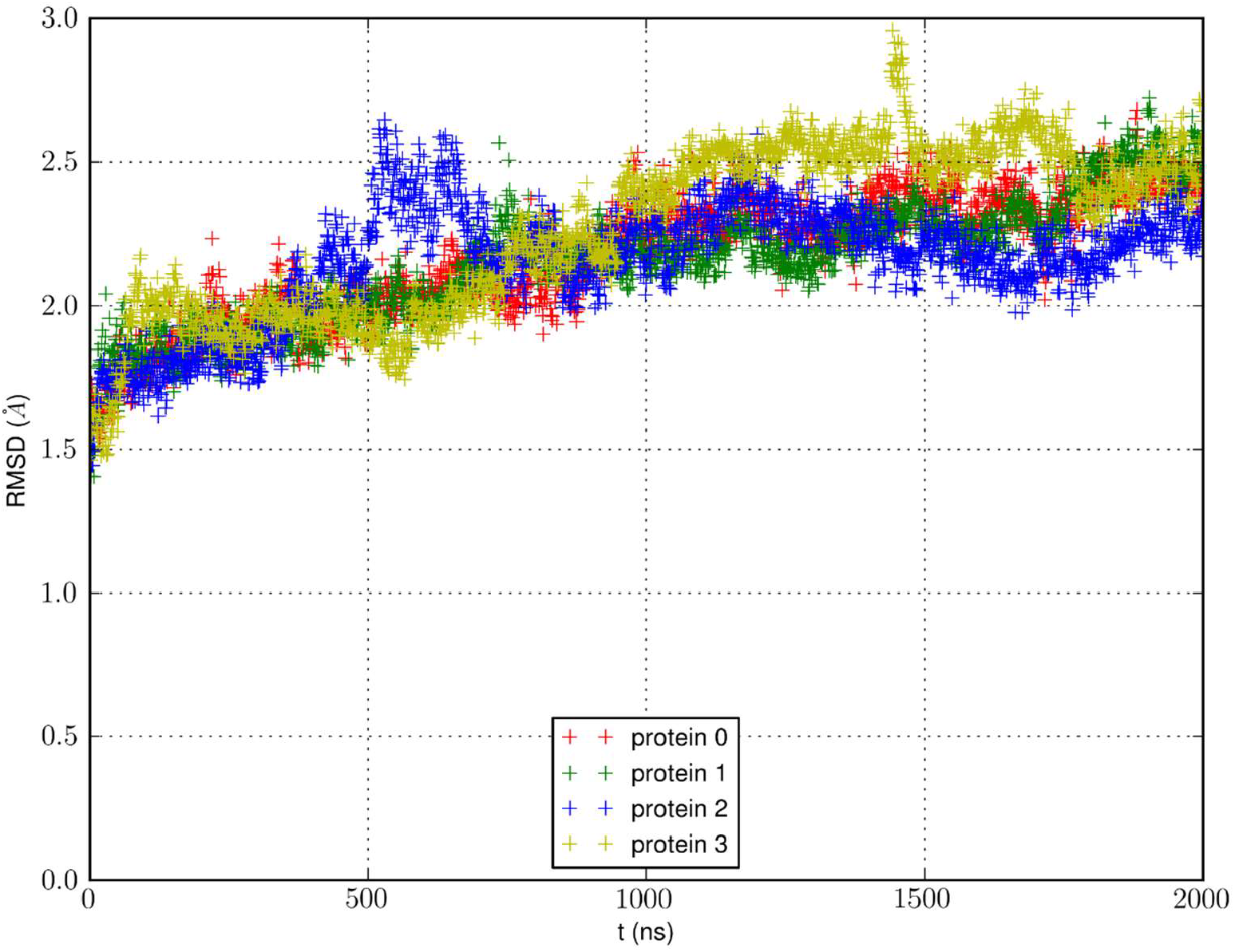
RMSD from the crystal structure of the protein monomers during the MD run of a system with a single AQP7 tetramer (4 monomer channel proteins) for 2,000 ns (*i*.*e*., 8,000 monomer•ns).

### Small simulations suggest low glycerol-AQP7 affinity

Analyzing the MD trajectory of SysI, a system consisting of one AQP7 tetramer in the presence of 50 mM glycerol, we counted one or more glycerol molecules residing inside a monomer channel as glycerol being within 7.1 Å from the NAA/NPS motifs located in the central part of the AQP7 channel. The probability of a channel being occupied by one or more glycerol molecules is shown in Fig. 4 along with the probability of a channel being occupied by two glycerol molecules. From the last 500 ns (2,000 monomer•ns), we computed the probability of an AQP7 channel being occupied by glycerol 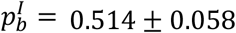 leading to a computed value of the glycerol-AQP7 dissociation constant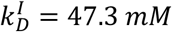. The computed affinity is not high, far from the experimentally measured value of 0.01 *mM* [7].

**Fig 4.**
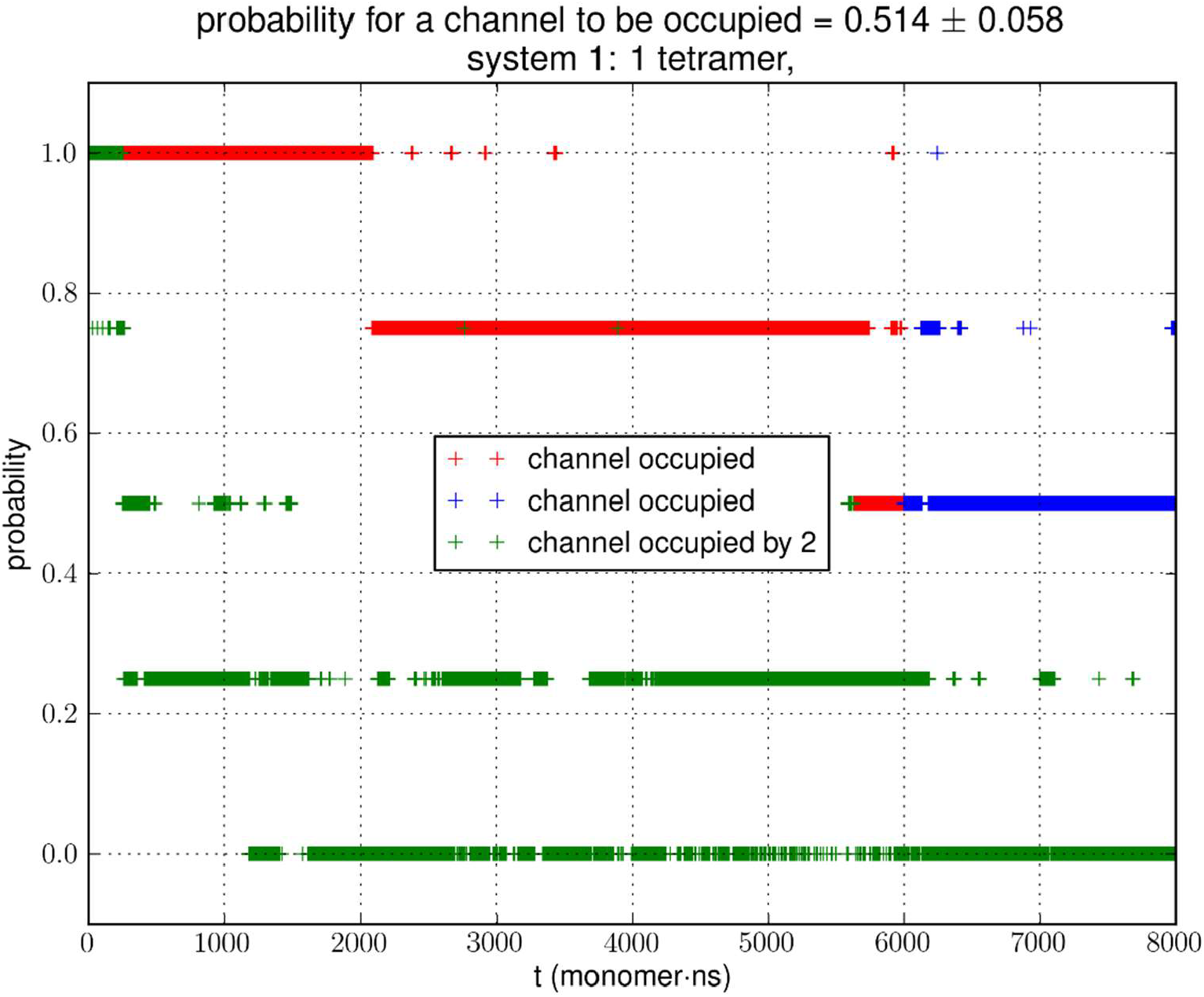
Glycerol binding characteristics of one tetramer in a typical simulation. The last 500 ns of the trajectory (*i*.*e*., 2,000 monomer•ns of dynamics, colored in blue) was used in the statistical calculation of the probability. During the first 1,500 ns shown in red, the system has not reached equilibrium. A channel is considered occupied where one or more glycerol molecules are within 7.1 Å from the NAA/NPS motifs.

Does the large discrepancy from the *in vitro* data mean that *in silico* studies cannot be quantitatively accurate at all? Where does this large discrepancy come from? Our model system (SysI) is typical of the current literature[15]. We used the standard CHARMM force field parameters. We did not employ any biases in the MD simulation that can generated artifacts. However, during the NPT run for a constant pressure of 1.0 bar, the model system was actually subjected to the mechanic pressure that fluctuated between ±400 bar (Fig. 1). This pressure fluctuation is inevitable in any simulations because it is intrinsic to any system that is smaller than the thermodynamic limit. The mean square fluctuation of pressure is inversely proportional to the system volume (thus the number of atoms constituting the model system) [21]. In light of all this, it is only logical to build larger systems to ascertain whether or not the pressure fluctuations caused the glycerol-AQP7 affinity to appear weak.

### Larger simulations yield greater estimates of the glycerol-AQP7 affinity

In Fig. 5, we show the results of 15,000 monomer•ns simulations of two larger systems, SysII consisting of 4.3 M atoms and SysIII consisting of 10.2 M atoms. The pressure fluctuations of these two systems (shown in Fig. 1) are significantly smaller than SysI. The mean square fluctuations are approximately in inverse proportion to the system size (the number of atoms) as expected from statistical thermodynamics[21]. Using the same criterium as for SysI, when one or more glycerol molecules are within 7.1 Å of the NAA/NPS motifs of an AQP7 monomer, that AQP7 channel is counted as being occupied. For a given time interval, SysII and SysIII have many more events of glycerol binding to and dissociating from an AQP7 channel than SysI. Naturally, with larger simulations, we have better statistics in addition to the fact that we have much smaller artifactitious fluctuations in pressure. Taking the last 5,000 monomer•ns of the MD trajectories into the statistical calculations, we obtained the probability for an AQP7 channel being occupied by glycerol, 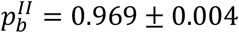for SysII and 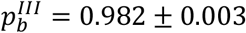 for SysIII. Correspondingly, the computed values of glycerol-AQP7 dissociation constant are 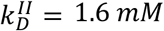 for SysII and 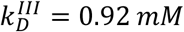for SysIII. Considering the computed value for SysI, 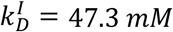, we observe the convergence toward higher affinities (lower *k*_D_ values) in larger model systems. There is a strong correlation between the computed *k*_D_ values and the artifactitious pressure fluctuations that are inevitable in any computational studies. Ideally, one can build a large enough system whose pressure fluctuation is much less than 1.0 bar for NPT runs under a constant pressure of 1.0 bar, which is still infeasible with today’s computing power. However, our study of SysI, SysII, and SysIII together showed that the glycerol-AQP7 affinity is indeed high as one would expect for a facilitator protein with its substrate. It is emphasized here that the afore-presented computations are directly from unbiased equilibrium MD simulations. As long as the parameters are accurate for the intra- and inter-molecular interactions, the conclusion of high glycerol-AQP7 affinity should be valid, free from artifacts that may be present in biased MD simulations.

**Fig 5.**
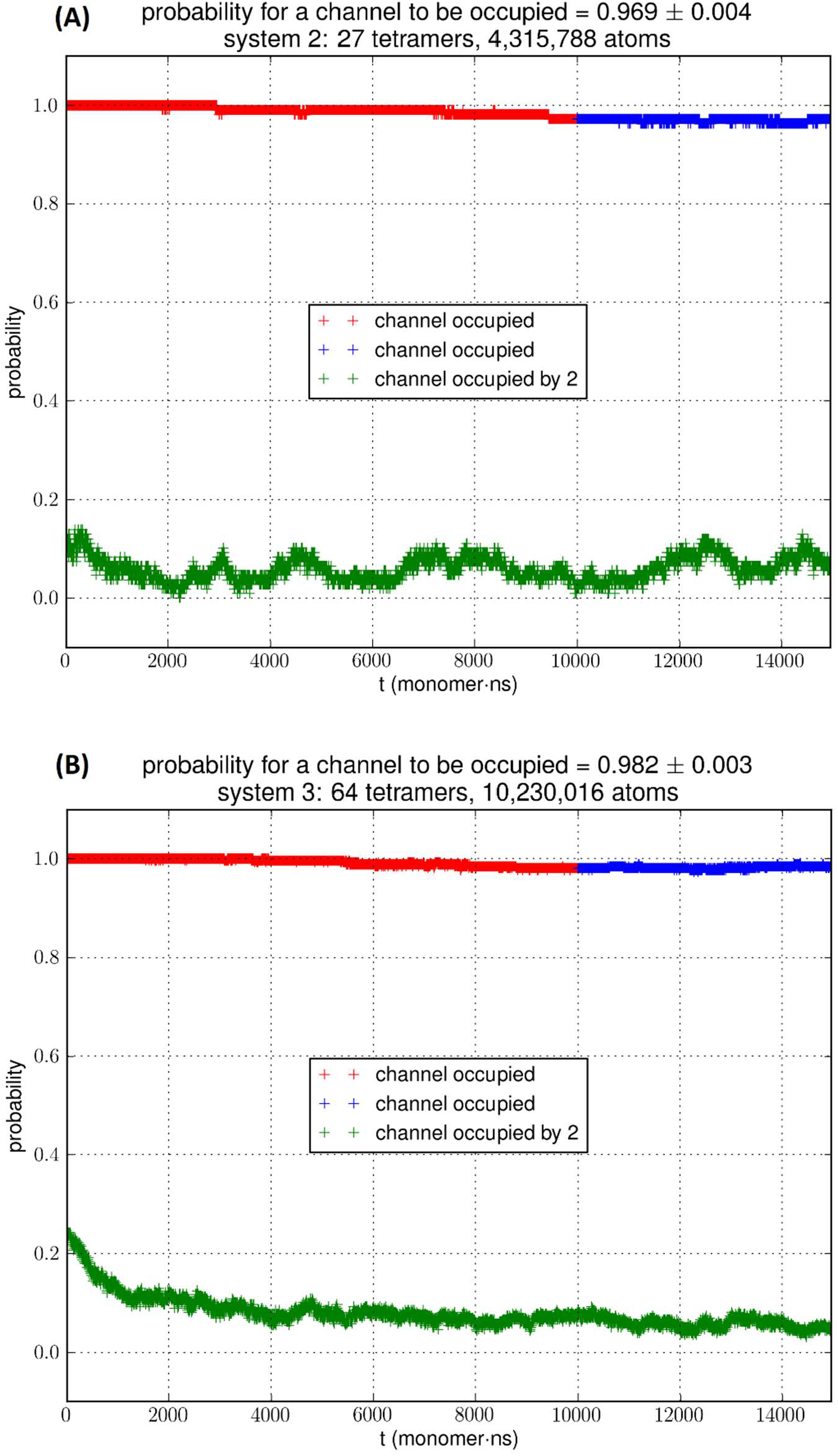
**(A)** SysII, 27 tetramers in a large simulation. **(B)** SysIII, 64 tetramers in a huge simulation. The last 5,000 monomer·ns of the trajectory (colored in blue) was used in the statistical calculation of the probability. More details are shown SI, Figs. S4 and S5.

### Affinity from the Gibbs free-energy profile

Fig. 6 shows the PMF throughout the AQP7 channel as a function of the z-coordinate of the glycerol’s center of mass. The PMF was computed from hSMD sampling of glycerol transport through AQP7 illustrated in Movie 2. It represents the Gibbs free energy of the system when a glycerol molecule is located at a given location. The reference level of the PMF was chosen at the bulk level on either the EC or the IC side. The two bulk levels must be equal for neutral solute transport across the cell membrane which is not an actively driven process but a facilitated passive process of diffusion down the concentration gradient. The PMF curve leveling off to zero on both the EC side and the IC side in Fig. 6 indicates accuracy of our computation. Inside the protein channel, the PMF presents a deep well (Δ*W*_0_ = −9.2 kcal/mol) near the NAA/NPS motifs (around *z*∼ − 1), which is a binding site for glycerol (Site 0). On the EC side, near the aromatic/Arginine (ar/R) selectivity filter (sf), there is another binding site (Site 1) where the PMF has a local minimum (Δ*W*_l_ = −4.7 kcal/mol). The third binding site (Site 2) is located on the IC side of the NAA/NPS where the PMF is Δ*W*_2_ = −3.3 kcal/mol. The PMF well depth is the main factor to determine the affinity (the inverse dissociation constant) at a given binding site. 1/*k*_D_ = *f*_O_exp [−Δ*W*_0_/*RT*] for the central binding site. The other factors involved in the determination of the affinities are the fluctuations (shown in SI, Figs. S7 to S10) which were computed straightforwardly from the equilibrium MD runs with the gaussian approximation. Combining the fluctuations and the PMF well depth, we obtained the dissociation constants as follows: *k*_D_ = 0.18 *mM* for the central binding site. This independent computation of the AQP7-glycerol affinity from hSMD simulations supports our direct computation from equilibrium MD simulations (*k*_D_ < 0.92 *mM*).

**Fig. 6.**
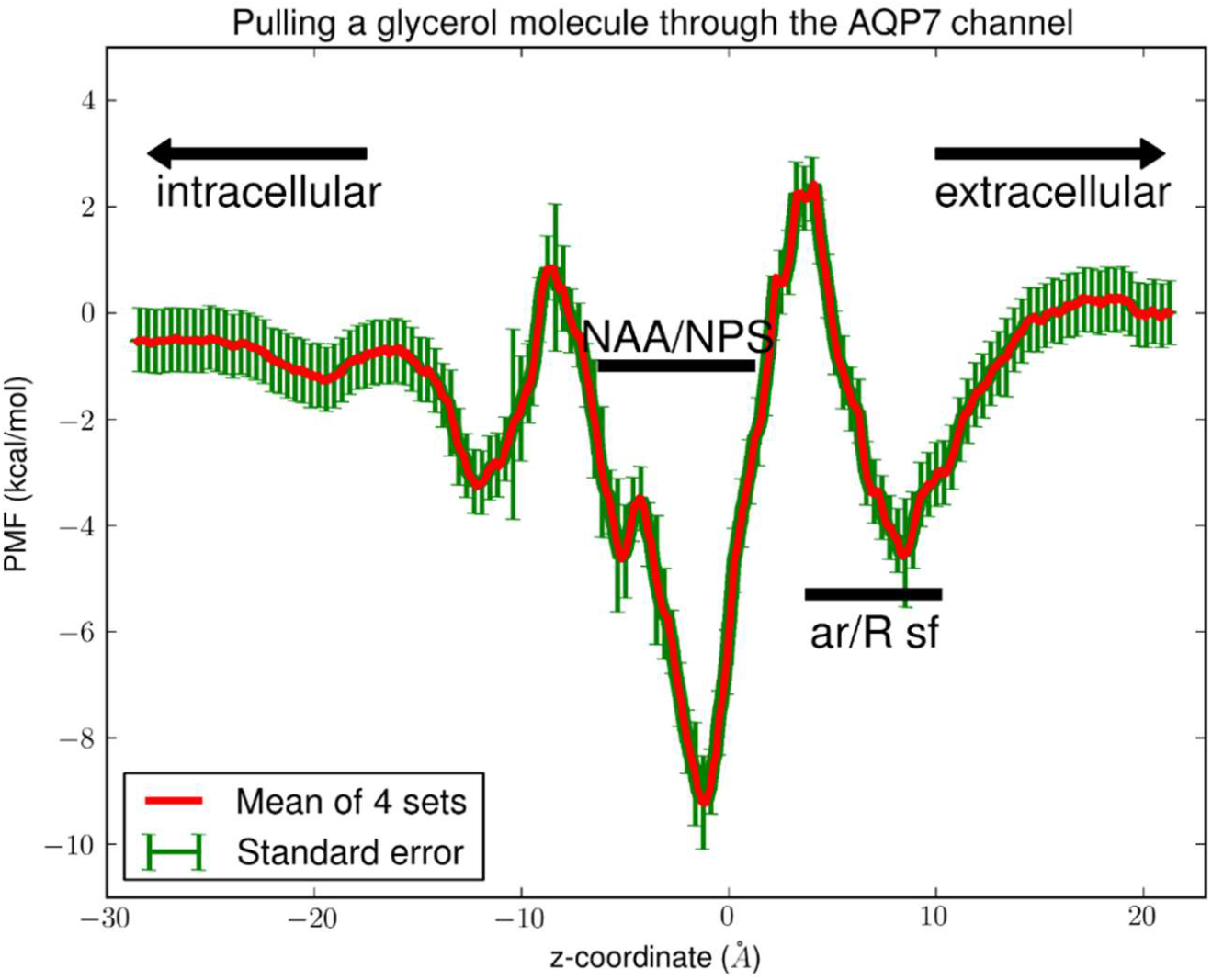
PMF of glycerol throughout the AQP7 channel. The coordinates are set so that the center of membrane is located at z∼0Å. In the single-file region (−11Å<z<9Å), the PMF is one dimensional. In the IC (z<-11Å) or EC (z>9Å) side of the channel, the PMF is three dimensional. The three PMF wells (binding sites) are located at: Site 0, z∼-1Å; Site 1, z∼9Å; Site 2, z∼-11Å.

## CONCLUSIONS

Based on the unbiased MD simulations of typically sized system and two very large systems, we observed that larger pressure fluctuations in smaller systems cause the glycerol-aquaglyceroporin affinity to appear lower. Beyond the consequence of the artifactitious pressure fluctuations, the computed values of glycerol-AQP7 dissociation constant indicate high affinity of an aquaglyceroporin for its substrate, which is in agreement with the *in vitro* data on AQP7.

## Supplementary Information

Two movies and 10 additional figures.

## Data availability

The Dataset (parameters, coordinates, scripts, *etc*.) to replicate this study is available at Harvard Dataverse[22].

## Author contributions

MF and LYC did the computational work; LYC conceptualized the research and wrote the paper; All participated in analyzing the data and editing the manuscript.

## Declaration

There are no conflicts to declare.

## Acknowledgements

The authors acknowledge use of computational time on Frontera at the Texas Advanced Computing Center at the University of Texas at Austin. Frontera is made possible by National Science Foundation award OAC-1818253.

## REFERENCES

[1] P. Agre, L.S. King, M. Yasui, W.B. Guggino, O.P. Ottersen, Y. Fujiyoshi, A. Engel, S. Nielsen, Aquaporin water channels – from atomic structure to clinical medicine, The Journal of Physiology, 542 (2002) 3–16.

[2] G. Benga, On the definition, nomenclature and classification of water channel proteins (aquaporins and relatives), Molecular Aspects of Medicine, 33 (2012) 514–517.

[3] P. Agre, M. Bonhivers, M.J. Borgnia, The Aquaporins, Blueprints for Cellular Plumbing Systems, J. Biol. Chem., 273 (1998) 14659–14662.

[4] A. Engel, H. Stahlberg, Aquaglyceroporins: Channel proteins with a conserved core, multiple functions, and variable surfaces, in: W.D.S. Thomas Zeuthen (Ed.) International Review of Cytology, vol. 215, Academic Press, Cambridge, MA, 2002, pp. 75–104.

[5] G. Calamita, J. Perret, C. Delporte, Aquaglyceroporins: Drug Targets for Metabolic Diseases?, Frontiers in Physiology, 9 (2018) 851.

[6] C. Maurel, J. Reizer, J.I. Schroeder, M.J. Chrispeels, M.H. Saier, Functional characterization of the Escherichia coli glycerol facilitator, GlpF, in Xenopus oocytes, J. Biol. Chem., 269 (1994) 11869–11872.

[7] T. Katano, Y. Ito, K. Ohta, T. Yasujima, K. Inoue, H. Yuasa, Functional Characteristics of Aquaporin 7 as a Facilitative Glycerol Carrier, Drug Metabolism and Pharmacokinetics, 29 (2014) 244–248.

[8] M. Ishii, K. Ohta, T. Katano, K. Urano, J. Watanabe, A. Miyamoto, K. Inoue, H. Yuasa, Dual Functional Characteristic of Human Aquaporin 10 for Solute Transport, Cell. Physiol. Biochem., 27 (2011) 749–756.

[9] Y. Ohgusu, K.-y. Ohta, M. Ishii, T. Katano, K. Urano, J. Watanabe, K. Inoue, H. Yuasa, Functional Characterization of Human Aquaporin 9 as a Facilitative Glycerol Carrier, Drug Metabolism and Pharmacokinetics, 23 (2008) 279–284.

[10] D. Fu, A. Libson, L.J.W. Miercke, C. Weitzman, P. Nollert, J. Krucinski, R.M. Stroud, Structure of a Glycerol-Conducting Channel and the Basis for Its Selectivity, Science, 290 (2000) 481–486.

[11] Z.E. Newby, J. O’Connell, 3rd, Y. Robles-Colmenares, S. Khademi, L.J. Miercke, R.M. Stroud, Crystal structure of the aquaglyceroporin PfAQP from the malarial parasite Plasmodium falciparum, Nat Struct Mol Biol, 15 (2008) 619–625.

[12] K. Gotfryd, A.F. Mósca, J.W. Missel, S.F. Truelsen, K. Wang, M. Spulber, S. Krabbe,C. Hélix-Nielsen, U. Laforenza, G. Soveral, P.A. Pedersen, P. Gourdon, Human adipose glycerol flux is regulated by a pH gate in AQP10, Nature Communications, 9 (2018) 4749.

[13] S.W. de Maré, R. Venskutonytė, S. Eltschkner, B.L. de Groot, K. Lindkvist-Petersson, Structural Basis for Glycerol Efflux and Selectivity of Human Aquaporin 7, Structure, 28 (2020) 215-222.e213.

[14] L. Zhang, D. Yao, Y. Xia, F. Zhou, Q. Zhang, Q. Wang, A. Qin, J. Zhao, D. Li, Y. Li, L. Zhou, Y. Cao, The structural basis for glycerol permeation by human AQP7, Science Bulletin, 66 (2021) 1550–1558.

[15] F.J. Moss, P. Mahinthichaichan, D.T. Lodowski, T. Kowatz, E. Tajkhorshid, A. Engel, W.F. Boron, A. Vahedi-Faridi, Aquaporin-7: A Dynamic Aquaglyceroporin With Greater Water and Glycerol Permeability Than Its Bacterial Homolog GlpF, Frontiers in Physiology, 11 (2020) 728.

[16] R.A. Rodriguez, R. Chan, H. Liang, L.Y. Chen, Quantitative study of unsaturated transport of glycerol through aquaglyceroporin that has high affinity for glycerol, RSC advances, 10 (2020) 34203–34214.

[17] N. Roudier, J.-M. Verbavatz, C. Maurel, P. Ripoche, F. Tacnet, Evidence for the Presence of Aquaporin-3 in Human Red Blood Cells, J. Biol. Chem., 273 (1998) 8407–8412.

[18] M.Ø. Jensen, S. Park, E. Tajkhorshid, K. Schulten, Energetics of glycerol conduction through aquaglyceroporin GlpF, Proceedings of the National Academy of Sciences, 99 (2002) 6731–6736.

[19] J.S. Hub, B.L. de Groot, Mechanism of selectivity in aquaporins and aquaglyceroporins, Proc. Natl. Acad. Sci. U.S.A., 105 (2008) 1198–1203.

[20] L.Y. Chen, Glycerol modulates water permeation through Escherichia coli aquaglyceroporin GlpF, Biochimica et Biophysica Acta (BBA) - Biomembranes, 1828 (2013) 1786–1793.

[21] L.D. Landau, E.M. Lifshitz, Statistical Physics, Part 1, Pergamon Press, Tarrytown, 1985.

[22] L.Y. Chen, Replication Data for “Aquaglyceroporin AQP7 s affinity for its substrate glycerol---Have we reached convergence in the computed values of glycerol-aquaglyceroporin affinity?”, in, Harvard Dataverse, 2021.

[23] S. Jo, T. Kim, V.G. Iyer, W. Im, CHARMM-GUI: A web-based graphical user interface for CHARMM, J. Comput. Chem., 29 (2008) 1859–1865.

[24] J. Lee, X. Cheng, J.M. Swails, M.S. Yeom, P.K. Eastman, J.A. Lemkul, S. Wei, J. Buckner, J.C. Jeong, Y. Qi, S. Jo, V.S. Pande, D.A. Case, C.L. Brooks, A.D. MacKerell, J.B. Klauda, W. Im, CHARMM-GUI Input Generator for NAMD, GROMACS, AMBER, OpenMM, and CHARMM/OpenMM Simulations Using the CHARMM36 Additive Force Field, Journal of Chemical Theory and Computation, 12 (2016) 405–413.

[25] J. Lee, D.S. Patel, J. Ståhle, S.-J. Park, N.R. Kern, S. Kim, J. Lee, X. Cheng, M.A. Valvano, O. Holst, Y.A. Knirel, Y. Qi, S. Jo, J.B. Klauda, G. Widmalm, W. Im, CHARMM-GUI Membrane Builder for Complex Biological Membrane Simulations with Glycolipids and Lipoglycans, Journal of Chemical Theory and Computation, 15 (2019) 775–786.

[26] J.C. Phillips, R. Braun, W. Wang, J. Gumbart, E. Tajkhorshid, E. Villa, C. Chipot, R.D. Skeel, L. Kalé, K. Schulten, Scalable molecular dynamics with NAMD, J. Comput. Chem., 26 (2005) 1781–1802.

[27] J.C. Phillips, D.J. Hardy, J.D.C. Maia, J.E. Stone, J.V. Ribeiro, R.C. Bernardi, R. Buch, G. Fiorin, J. Hénin, W. Jiang, R. McGreevy, M.C.R. Melo, B.K. Radak, R.D. Skeel, A. Singharoy, Y. Wang, B. Roux, A. Aksimentiev, Z. Luthey-Schulten, L.V. Kalé, K. Schulten, C. Chipot, E. Tajkhorshid, Scalable molecular dynamics on CPU and GPU architectures with NAMD, The Journal of Chemical Physics, 153 (2020) 044130.

[28] R.B. Best, X. Zhu, J. Shim, P.E.M. Lopes, J. Mittal, M. Feig, A.D. MacKerell, Optimization of the Additive CHARMM All-Atom Protein Force Field Targeting Improved Sampling of the Backbone φ, ψ and Side-Chain χ1 and χ2 Dihedral Angles, Journal of Chemical Theory and Computation, 8 (2012) 3257–3273.

[29] K. Vanommeslaeghe, E. Hatcher, C. Acharya, S. Kundu, S. Zhong, J. Shim, E. Darian, O. Guvench, P. Lopes, I. Vorobyov, A.D. Mackerell, CHARMM general force field: A force field for drug-like molecules compatible with the CHARMM all-atom additive biological force fields, J. Comput. Chem., 31 (2010) 671–690.

[30] J.B. Klauda, R.M. Venable, J.A. Freites, J.W. O’Connor, D.J. Tobias, C. Mondragon-Ramirez, I. Vorobyov, A.D. MacKerell, R.W. Pastor, Update of the CHARMM All-Atom Additive Force Field for Lipids: Validation on Six Lipid Types, The Journal of Physical Chemistry B, 114 (2010) 7830–7843.

[31] S. Padhi, U.D. Priyakumar, Selectivity and transport in aquaporins from molecular simulation studies, Vitam Horm, 112 (2020) 47–70.

[32] L.S.M. Neumann, A.H.S. Dias, M.S. Skaf, Molecular Modeling of Aquaporins from Leishmania major, The Journal of Physical Chemistry B, 124 (2020) 5825–5836.

[33] J.A. Freites, K.L. Németh-Cahalan, J.E. Hall, D.J. Tobias, Cooperativity and allostery in aquaporin 0 regulation by Ca2+, Biochimica et Biophysica Acta (BBA) - Biomembranes, 1861 (2019) 988–996.

[34] L.Y. Chen, Exploring the free-energy landscapes of biological systems with steered molecular dynamics, PCCP, 13 (2011) 6176–6183.

[35] W. Humphrey, A. Dalke, K. Schulten, VMD: Visual molecular dynamics, Journal of Molecular Graphics, 14 (1996) 33–38.

[36] H.-J. Woo, B. Roux, Calculation of absolute protein–ligand binding free energy from computer simulations, Proc. Natl. Acad. Sci. U.S.A., 102 (2005) 6825–6830.

[37] R.A. Rodriguez, L. Yu, L.Y. Chen, Computing Protein-Protein Association Affinity with Hybrid Steered Molecular Dynamics, J Chem Theory Comput, 11 (2015) 4427–4438.

