## Supplementary material for "Aquaglyceroporin AQP7’s affinity for its substrate glycerol": movies and supplemental figures

### Have we reached convergence in the computed values of glycerol-aquaglyceroporin affinity?

Michael Falato[1], Ruth Chan[1], and Liao Y. Chen[1*]

[1]Department of Physics, University of Texas at San Antonio, San Antonio, Texas 78249 USA

In this supplemental information, we show two movies and 10 additional figures that are discussed but not included in the main text.

movie1_binding-dissociating.mp4

**Movie 1.** Glycerol dissociating from AQP7 and binding to AQP7. The membrane lipids are shown in lines colored by atoms (H, white; C, cyan; N, blue; O, red; P, purple). Water and salt atoms are not shown for clarity. Glycerols are shown in space-filling spheres colored by atoms (*ibid.*). The AQP7 tetramer is shown in cartoons colored by monomers (monomer 1, blue; 2, red; 3, gray; 4, gold).

movie2_transport.mp4

**Movie 2.** Transport path of glycerol through the AQP7 channel across the cell membrane. AQP7 is shown as cartoons colored by residue types (positively charged, blue; negatively charged, red; hydrophilic, green; hydrophobic, white). Glycerol and waters inside and near the channel are shown as shiny and shadowy space-filling spheres, respectively, all colored by atoms (H, white; C, cyan; O, red).

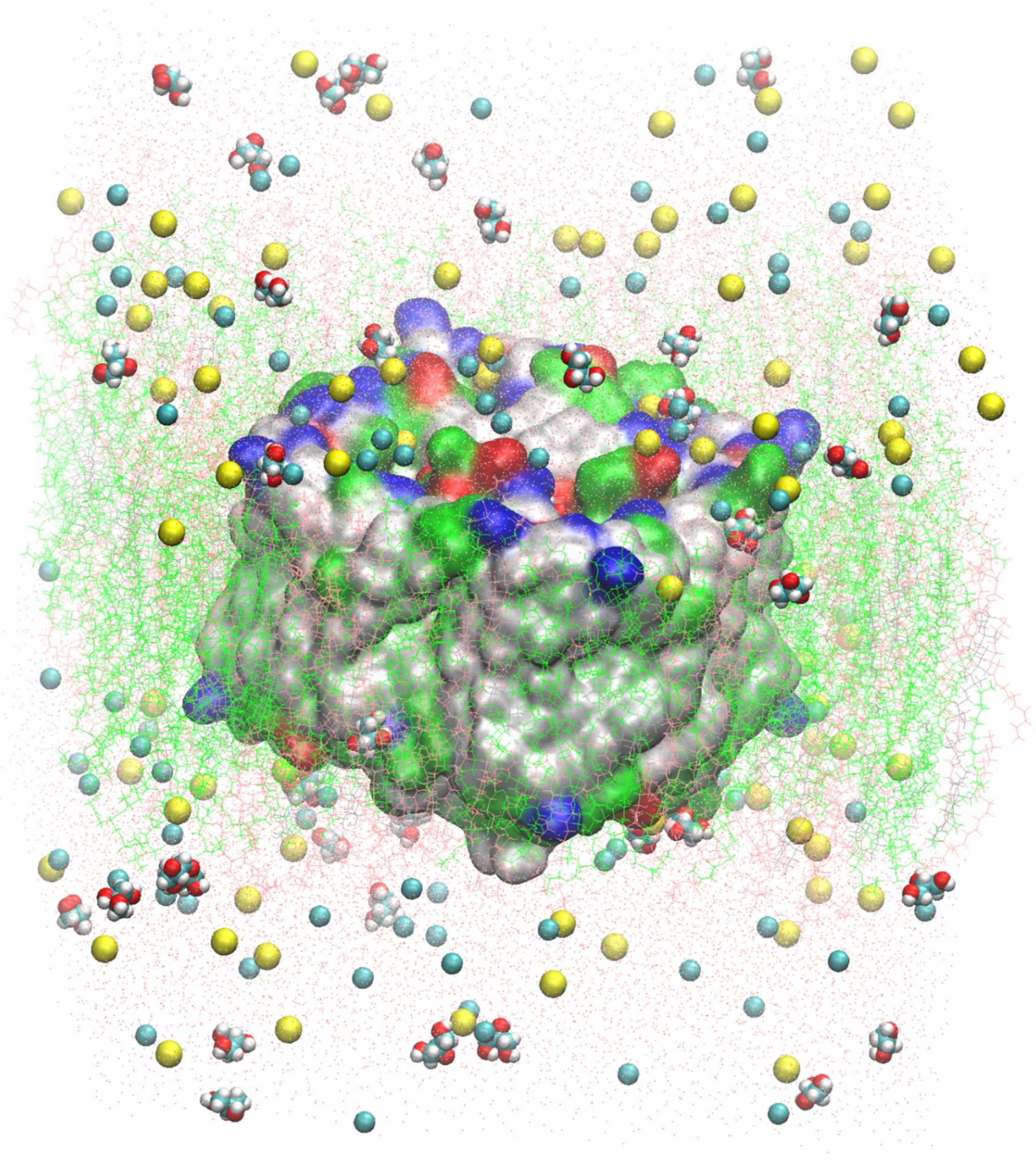

**Fig. S1.** System 1 (SysI, 159,844 atoms) consisting of one AQP7 tetramer embedded in a lipid bilayer solvated in 150 mM NaCl saline with 52 mM glycerol. The AQP7 tetramer is shown in surface colored by residue types (positively charged, blue; negatively charged, red; hydrophilic, green; hydrophobic, white). Membrane lipids are shown in lines colored by lipids (193 POPE's, green; 119 POPC's, pink; 80 Cholesterols, charcoal). Waters are shown in red dots. Glycerols and salt ions are shown in space-filling spheres colored by atoms (H, white; C, cyan; N, blue; O, red; Na, yellow; Cl, cyan).

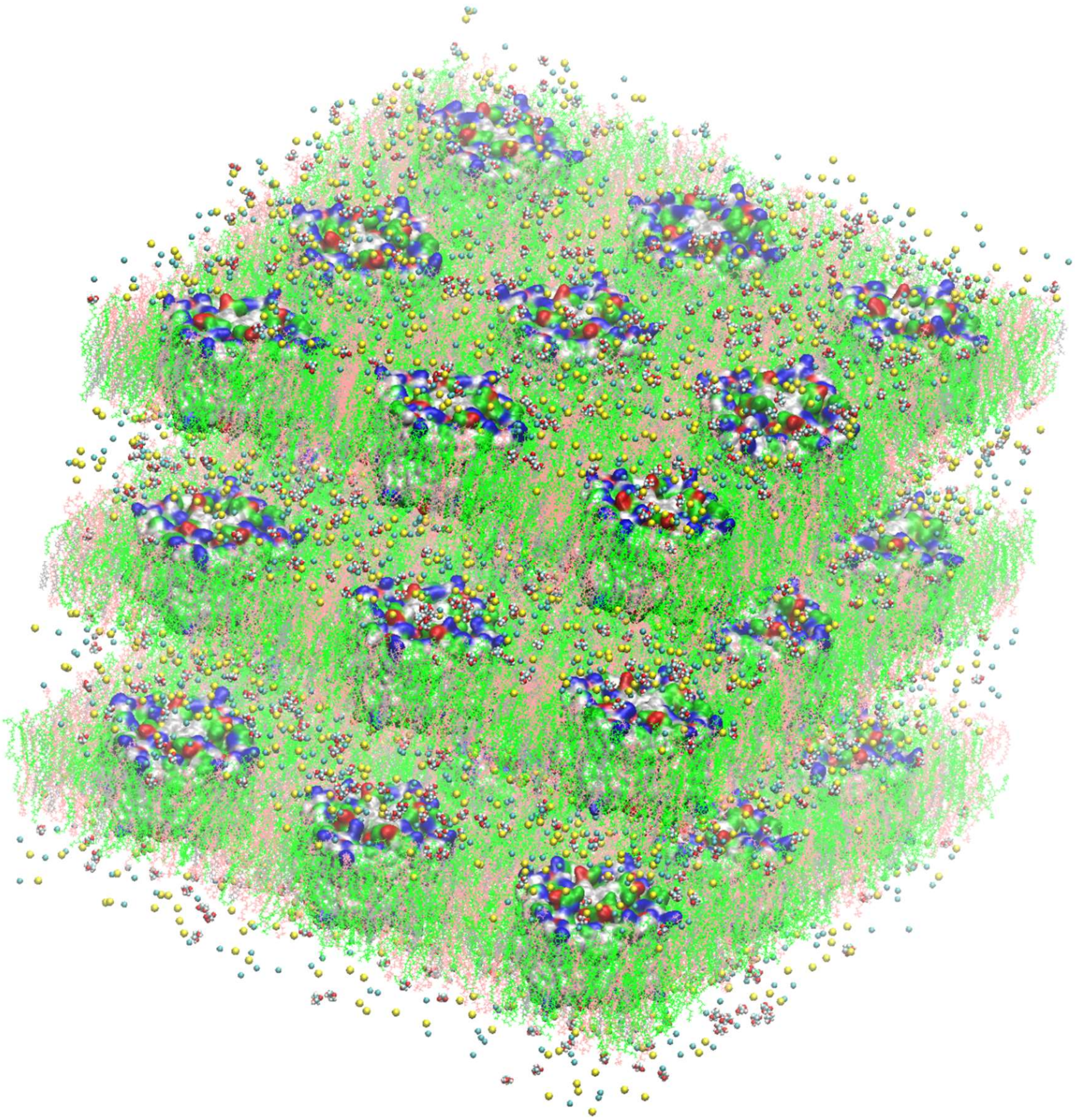

**Fig**. **S2**. System 2 (SysII, 4,313,628 atoms) consisting of 27 AQP7 tetramers embedded in three lipid bilayers solvated in 150 mM NaCl saline with 52 mM glycerol. Waters are shown for clarity. Representations and colors are identical to Fig. S1.

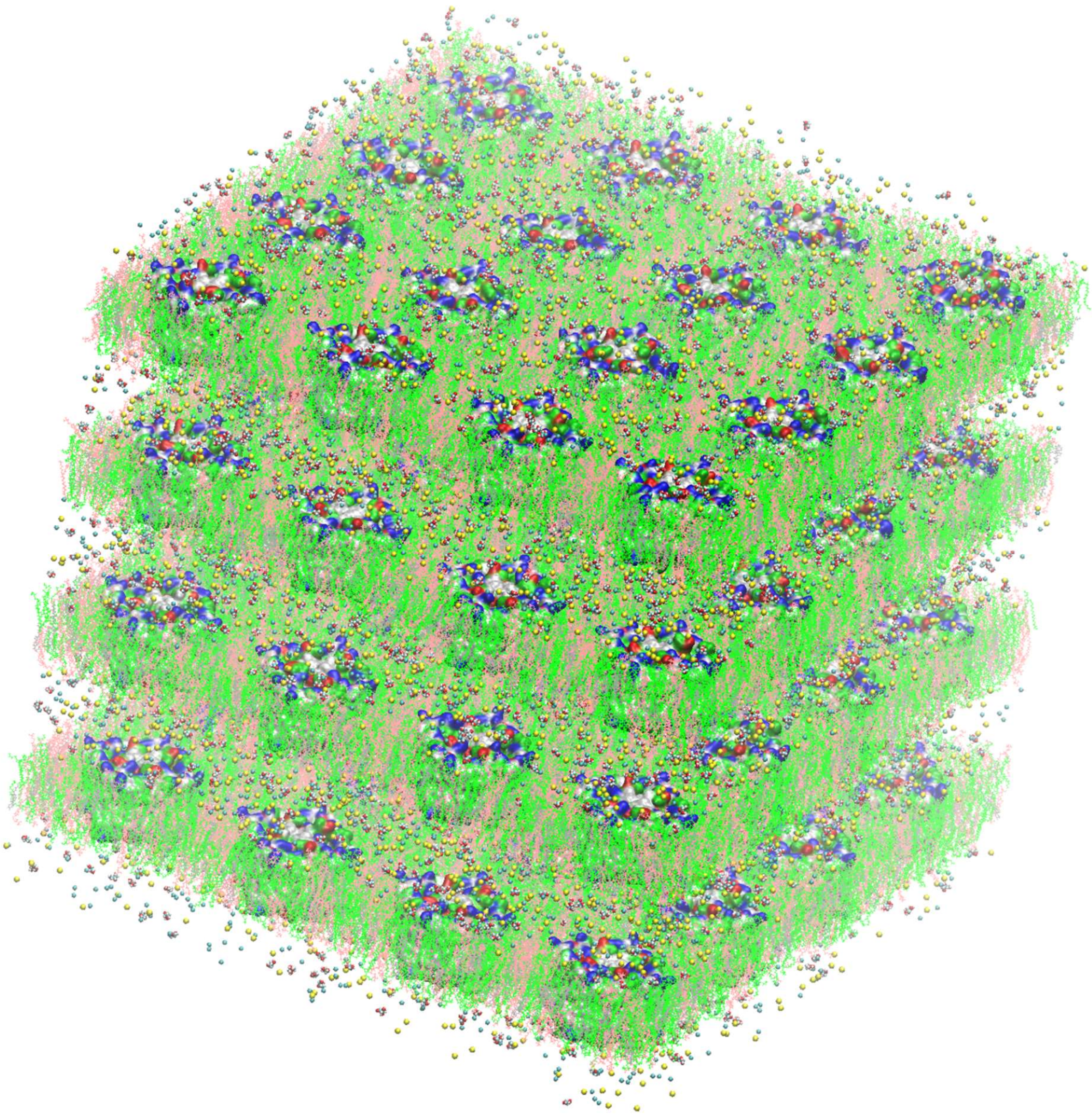

**Fig. S3.** System 3 (SysIII, 10,230,016 atoms) consisting of 64 AQP7 tetramers embedded in four lipid bilayers solvated in 150 mM NaCl saline with 52 mM glycerol. Waters are shown for clarity. Representations and colors are identical to Fig. S1.

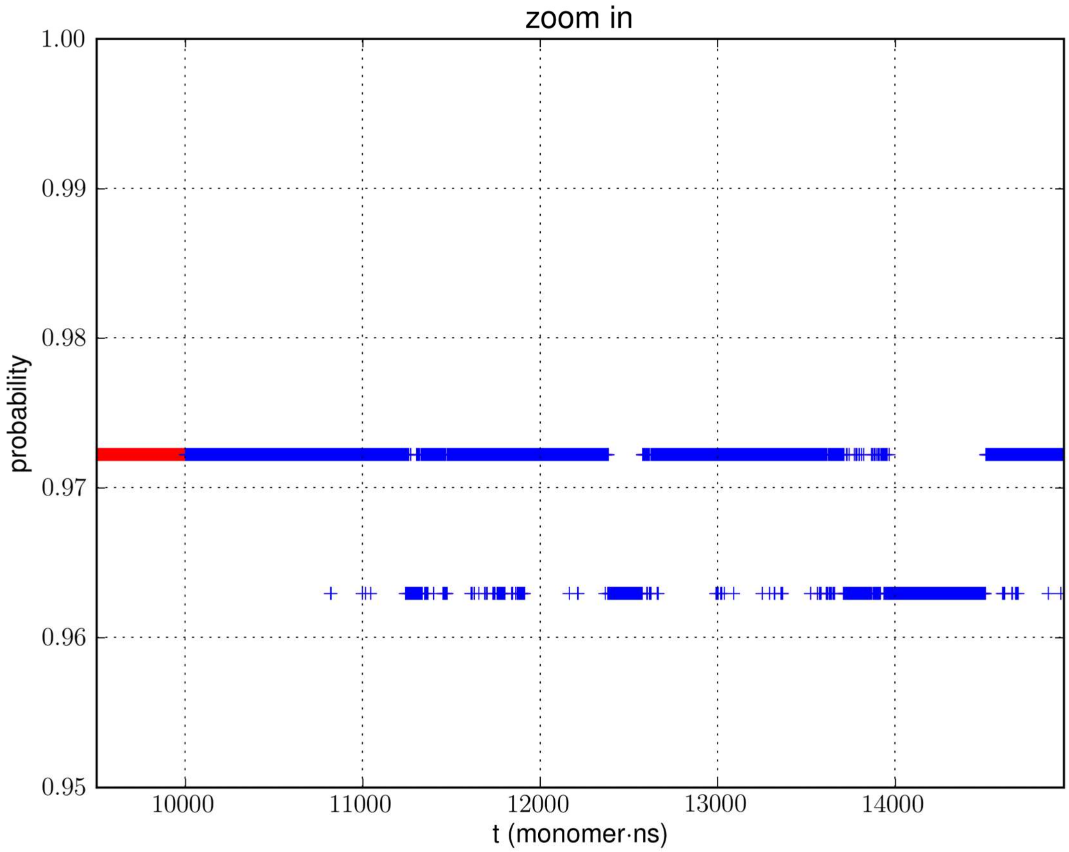


**Fig. S4.** Zoom-in details of the binding-dissociating activities in sysII, the system with 27 tetramers (108 monomer channel proteins).

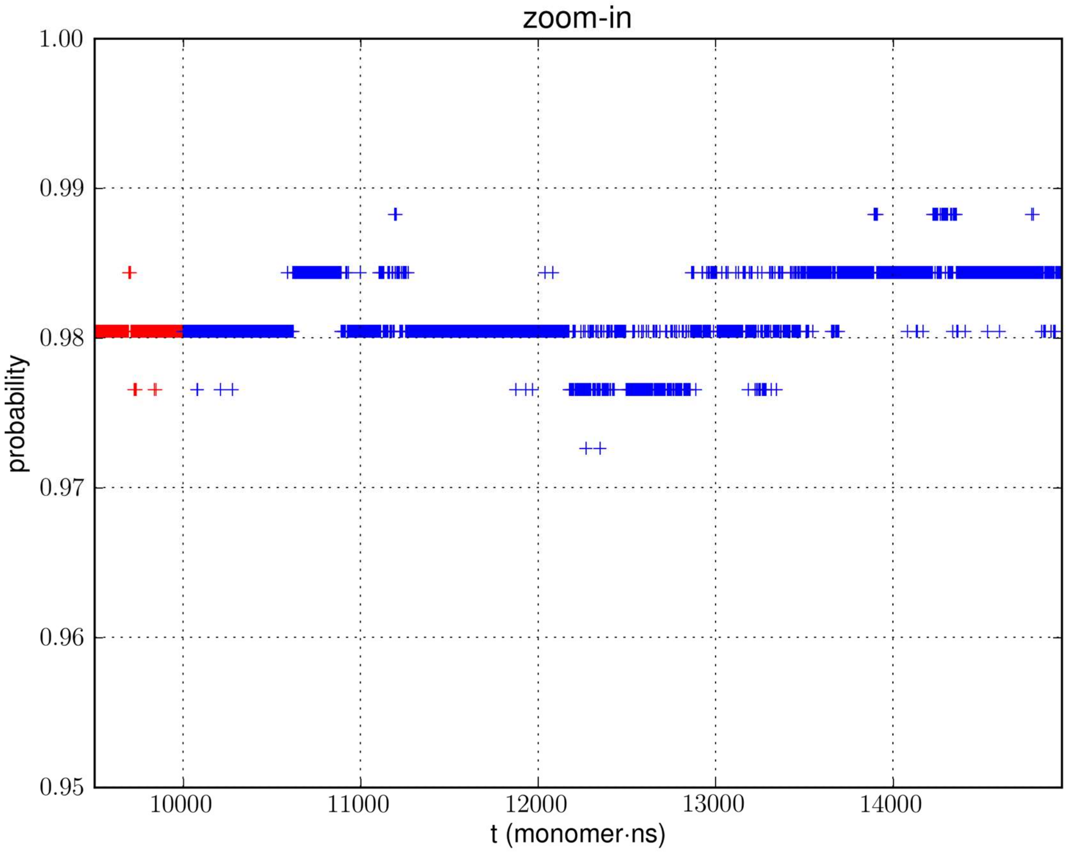


**Fig. S5.** Zoom-in details of the binding-dissociating activities in sysIII, the system with 64 tetramers (256 monomer channel proteins).

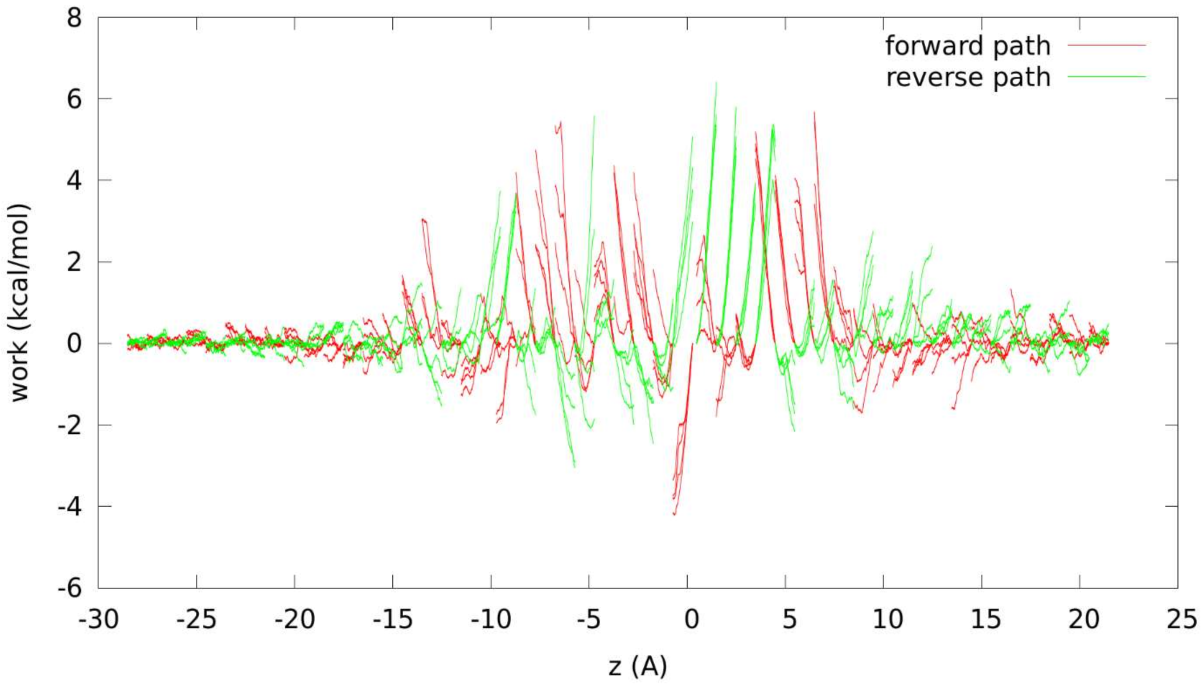


**Fig. S6.** Work done to the system when a glycerol molecule is hyper-steered along the forward and the reverse paths in each of the 50 sections from the IC side (z<-28), through the AQP7 channel, to the EC side (z>22) of the cell membrane.

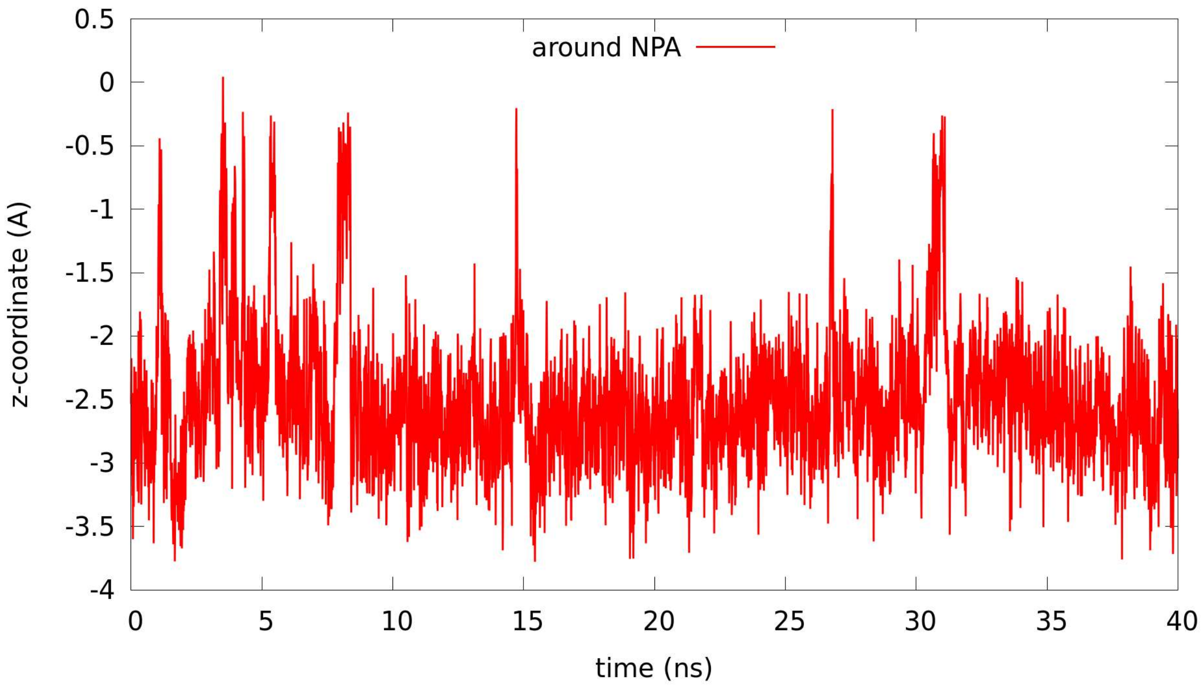


**Fig. S7**. Longitudinal (along the z-axis) fluctuations of glycerol at the central binding site around the "NPA" (NAA-NPS) motifs.

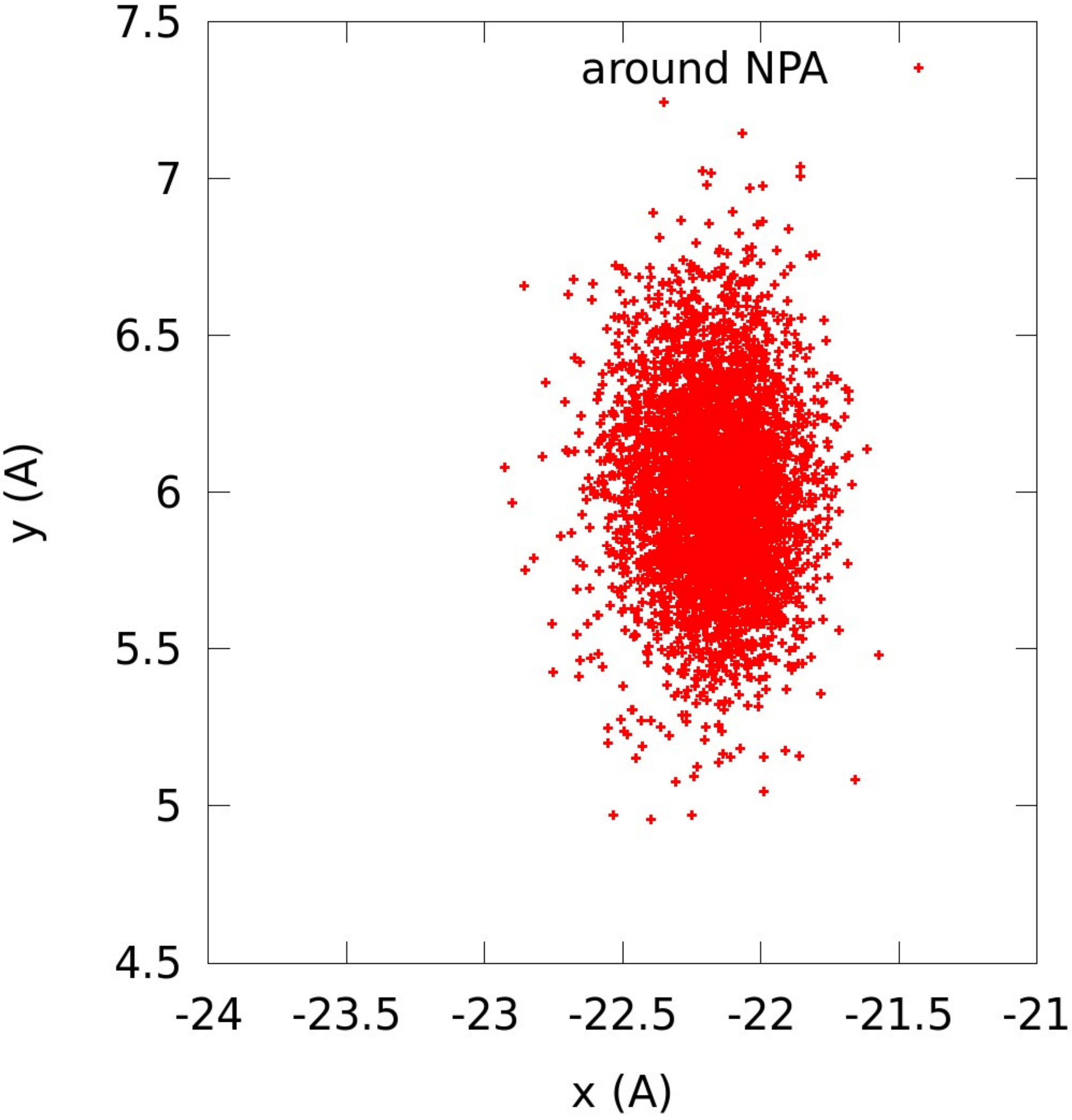


**Fig. S8**. Transverse fluctuations of glycerol at the central binding site around the “NPA” (NAA-NPS) motifs.

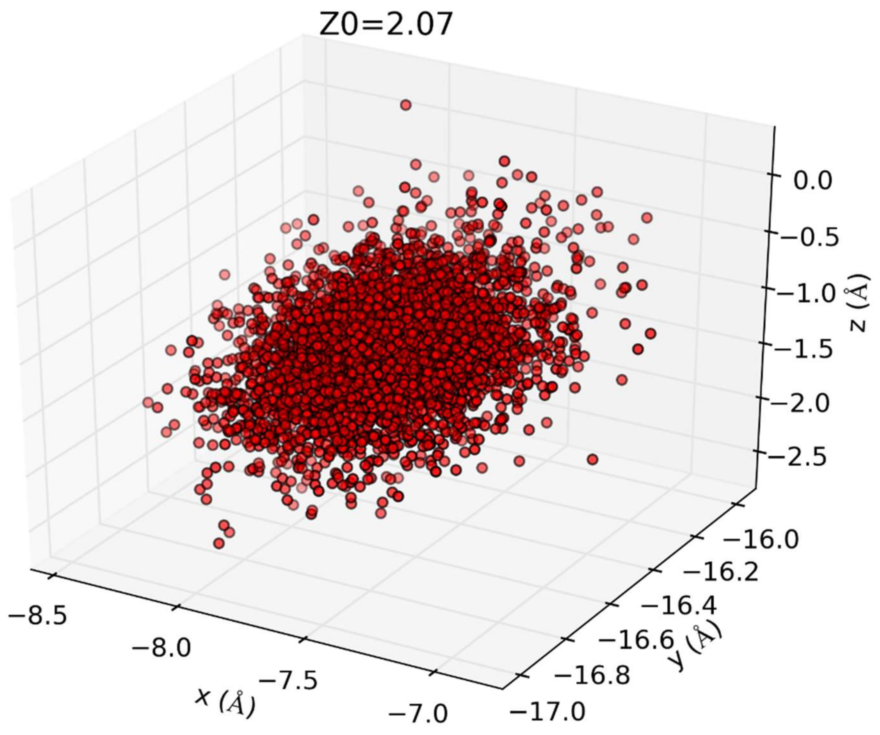


**Fig. S9**. Fluctuations of glycerol at the central binding site giving rise to the partial partition $Z_0$.

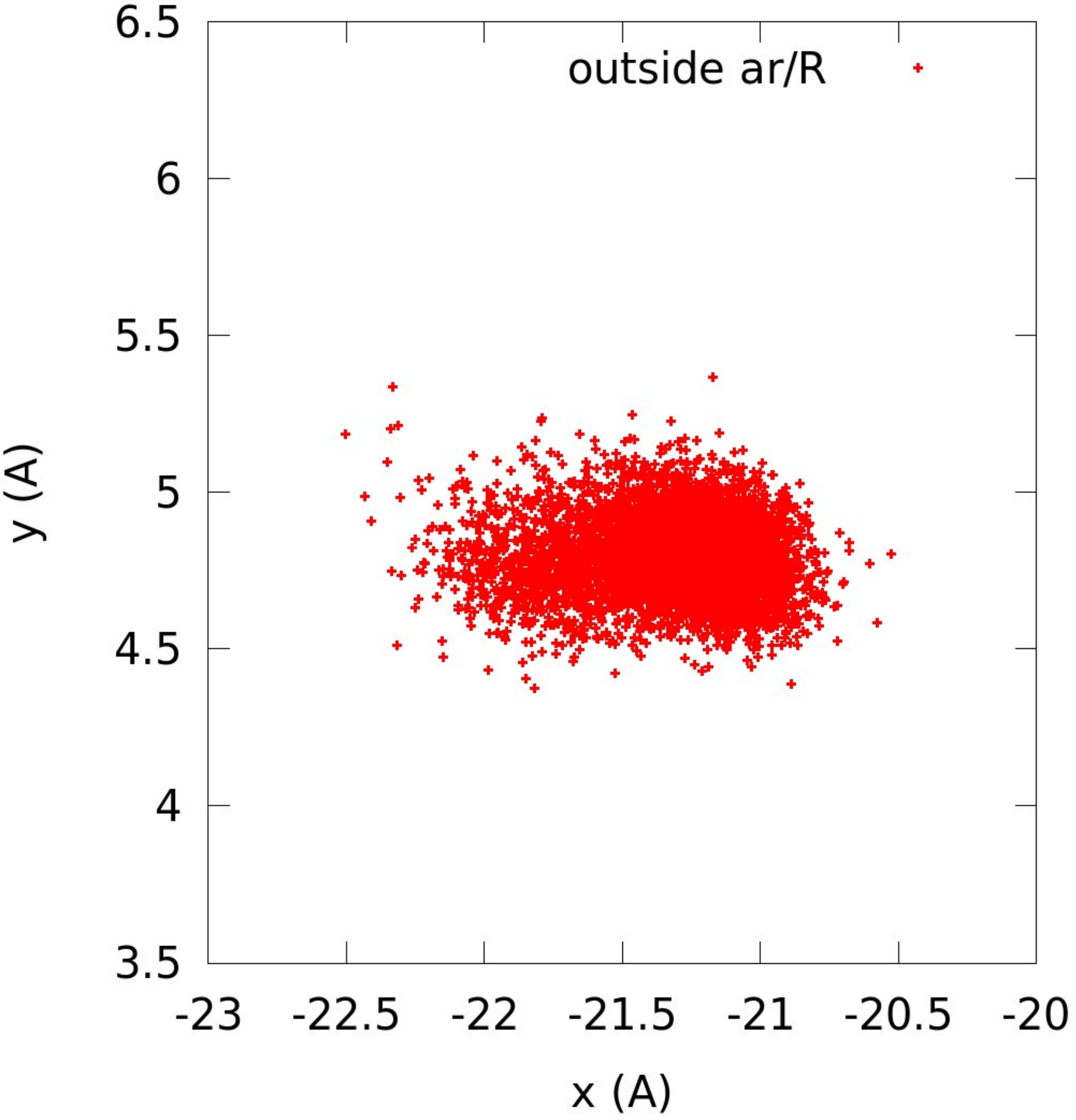


**Fig. S10**. Transverse fluctuations of glycerol at the channel opening on the EC side (a binding site near the ar/R).
